# Single-cell RNA-seq uncovers shared and distinct axes of variation in dorsal LGN neurons in mice, non-human primates and humans

**DOI:** 10.1101/2020.11.05.367482

**Authors:** Trygve E. Bakken, Cindy T.J. van Velthoven, Vilas Menon, Rebecca D. Hodge, Zizhen Yao, Thuc Nghi Nguyen, Lucas T. Graybuck, Gregory D. Horwitz, Darren Bertagnolli, Jeff Goldy, Emma Garren, Sheana Parry, Tamara Casper, Soraya I. Shehata, Eliza R. Barkan, Aaron Szafer, Boaz P. Levi, Nick Dee, Kimberly A. Smith, Susan M. Sunkin, Amy Bernard, John W. Phillips, Michael Hawrylycz, Christof Koch, Gabe Murphy, Ed Lein, Hongkui Zeng, Bosiljka Tasic

## Abstract

Abundant anatomical and physiological evidence supports the presence of at least three distinct types of relay glutamatergic neurons in the primate dorsal lateral geniculate nucleus (dLGN) of the thalamus, the brain region that conveys visual information from the retina to the primary visual cortex. Relay neuron diversity has also been described in the mouse dLGN (also known as LGd). Different types of relay neurons in mice, humans and macaques have distinct morphologies, distinct connectivity patterns, and convey different aspects of visual information to the cortex. To investigate the molecular underpinnings of these cell types, and how these relate to other cellular properties and differences in dLGN between human, macaque, and mice, we profiled gene expression in single nuclei and cells using RNA-sequencing. These efforts identified four distinct types of relay neurons in the primate dLGN, magnocellular neurons, parvocellular neurons, and two cell types expressing canonical marker genes for koniocellular neurons. Surprisingly, despite extensive documented morphological and physiological differences between magno- and parvocellular neurons, we identified few genes with significant differential expression between transcriptomic cell types corresponding to these two neuronal populations. We also detected strong donor-specific gene expression signatures in both macaque and human relay neurons. Likewise, the dominant feature of relay neurons of the adult mouse dLGN is high transcriptomic similarity, with an axis of heterogeneity that aligns with core vs. shell portions of mouse dLGN. Together, these data show that transcriptomic differences between principal cell types in the mature mammalian dLGN are subtle relative to striking differences in morphology and cortical projection targets. Finally, we align cellular expression profiles across species and find homologous types of relay neurons in macaque and human, and distinct relay neurons in mouse.

## INTRODUCTION

Relay neurons in the dorsal lateral geniculate nucleus (dLGN) of the thalamus are the primary conduit of information about visual stimuli detected by the retina to the primary visual cortex (Jones, 2007). In many mammals, including humans and non-human primates, relay neurons reside in three anatomically distinct sets of layers: the magnocellular, parvocellular, and koniocellular layers. The three groups of relay neurons represent a well-documented example of distinct cell types in the central nervous system; in the non-human primate and cat, these cells differ dramatically in size, receive input from different types of retinal ganglion cells (RGCs), innervate different layers of primary visual cortex, and respond preferentially to distinct features of visual stimuli (Livingstone and Hubel, 1988; Callaway, 2005). Magnocellular neurons have large somata, receive input from rod photoreceptors, have larger receptive field, provide higher contrast gain, and project to layer 4Cα of the primary visual cortex, whereas parvocellular neurons receive input from cone photoreceptors, have smaller receptive fields, lower contrast gain, and project to layer 4Cß of the primary visual cortex (Reid and Shapley, 2002; Jeffries et al., 2014). The cell bodies of koniocellular neurons reside in thin layers that are sandwiched between the magno- and/or parvocellular layers. At least some koniocellular neurons receive input from short-wavelength cones and project to layers 2-3 of the primary visual cortex (Hendry and Reid, 2000).

The mouse dLGN is not as clearly laminated as it is in most primates, but the cell bodies of its relay neurons can be divided into two subregions – core and shell – that receive different retinal input and project to different layers of visual cortex (Cruz-Martin et al., 2014). Shell neurons preferentially receive input from direction-selective retinal ganglion cells and project to layers 1-3 of the visual cortex, whereas core neurons mostly receive input from non-direction selective retinal ganglion cells and project to layer 4 (Cruz-Martin et al., 2014; Seabrook et al., 2017). In addition, mouse relay neurons have been classified based on their dendritic morphology into X, Y and W types (Krahe et al., 2011), although the correspondence between the projection target and local dendritic morphology has not been established.

Recent advances in single-cell and single-nucleus RNA-sequencing (scRNA-seq and snRNA-seq) provide a powerful, complementary approach to anatomical studies to delineate and distinguish types of neurons on the basis of their genome-wide gene expression profiles (Darmanis et al., 2015; Zeisel et al., 2015; Tasic et al., 2016; Saunders et al., 2018; Tasic et al., 2018; Zeisel et al., 2018; Hodge et al., 2019; Bakken et al., 2020). These techniques can be used to examine the conserved and unique features of cell types in different species (La Manno et al., 2016; Hodge et al., 2019; Bakken et al., 2020; Krienen et al., 2020). We profiled the dLGN of adult human, macaque, and mouse by scRNA-seq and snRNA-seq to identify transcriptomic cell types and investigate their correspondence across human, macaque, and mouse.

## RESULTS

### Single cell and nucleus transcriptomic profiling

We used our previously described experimental approach (Tasic et al., 2016; Bakken et al., 2018; Tasic et al., 2018) (**Fig. 1**, **Methods**) to isolate and transcriptomically profile nuclei from macaque (*Macaca nemestrina* and *Macaca fascicularis*) and human dLGN, as well as cells from mouse dLGN. Nuclei were collected from anatomically defined regions: magno- or parvo-cellular layers of non-human primate dLGN; konio-, magno- or parvo-cellular layers of human dLGN; and shell or core of mouse dLGN. Koniocellular layers are thin and dissections inevitably included cells from neighboring magno- and parvo-cellular layers. Adjacent thalamic nuclei were also sampled, including ventral pulvinar from a single macaque donor, and lateral posterior (LP) and ventral lateral geniculate (LGv) nuclei from several mice. Single cells or nuclei were isolated by FACS and enriched for neurons based on labeling with neuronal markers (NeuN in primates and tdTomato (tdT) in mouse). All single cells and nuclei were processed with SMART-seq v4 (Clontech) and Nextera XT (Illumina), and sequenced on HiSeq 2500 (Illumina). RNA-seq reads were aligned to corresponding genomes using the STAR aligner (Dobin et al., 2013), gene expression was quantified as the sum of intronic and exonic reads per gene and was normalized as counts per million (CPM) and log_2_-transformed as previously described (Tasic et al., 2018; Hodge et al., 2019). Seurat v3 R package was used for clustering (Butler et al., 2018; Stuart et al., 2019) (**Methods**). We report on 2003 macaque, 1209 human, and 2118 mouse QC-qualified single cell and nucleus transcriptomes with cluster-assigned identity (**Fig. S1A, Table S1**). Samples were sequenced to a median depth of 1.3 million reads/nucleus in macaque samples, 2.4 million reads/nucleus in human samples, and 2.5 million reads/cell for mouse samples (**Fig. S1B**). Median gene detection in macaque nuclei (~5,900) is lower than in human nuclei and mouse cells (~8,900 and ~9,000 respectively) (**Fig. S1C**). Macaque single nuclei were isolated from acute surgical tissue and based on our quality control during isolation as well as the clustering result (see below), we consider them to be at least as high quality as nuclei from post-mortem human tissue. The lower gene detection for macaque nuclei likely resulted from alignment to genome of a related species, rhesus macaque (*Macaca mulatta*; see **Methods**), which we chose to use as it is better annotated than genomes of *Macaca nemestrina* and *Macaca fascicularis*.

**Figure 1.**
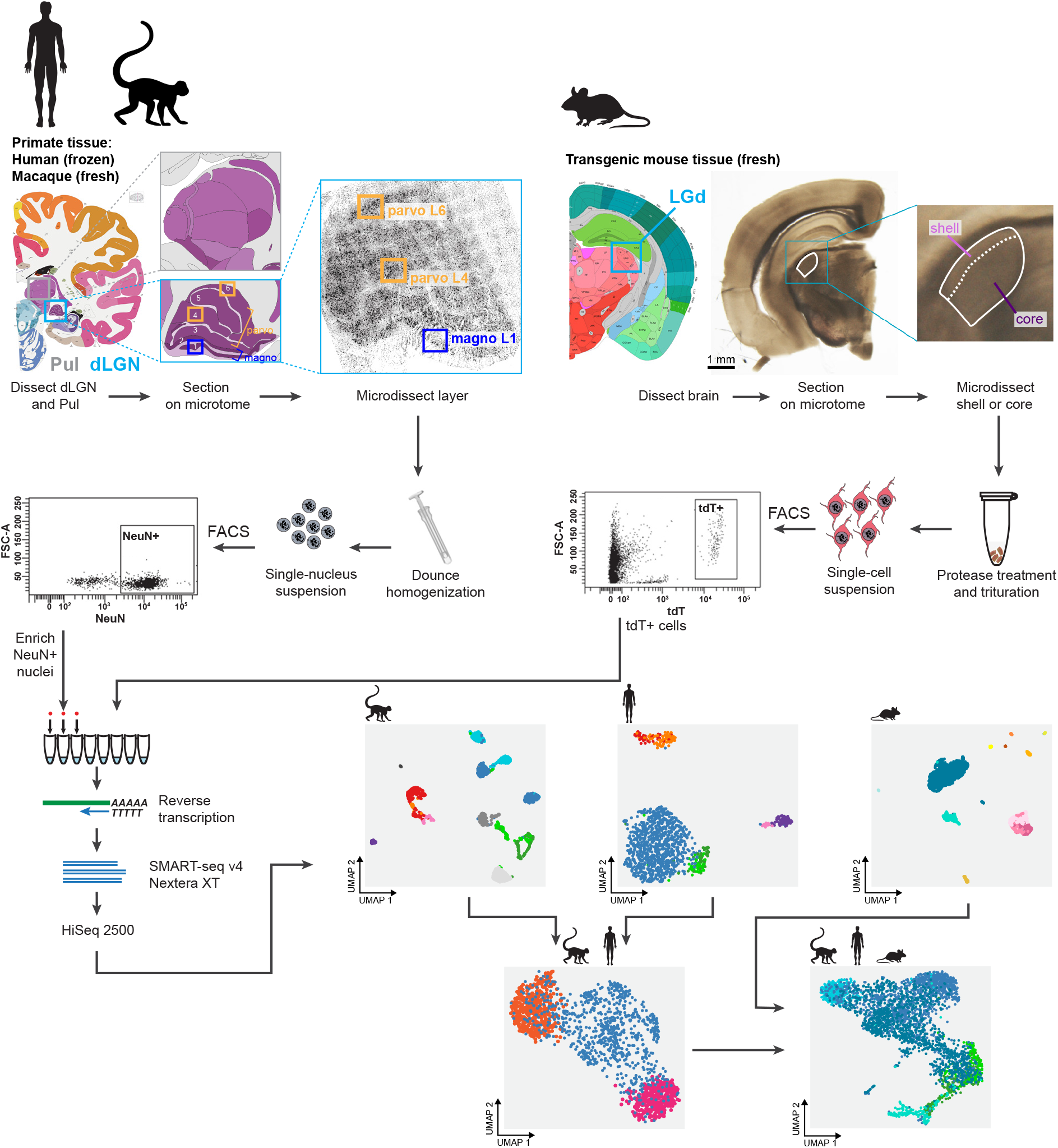
Experimental workflow and sample collection. dLGN was dissected from postmortem human brain, and acutely collected macaque and mouse brain according to the Allen Brain Atlas. Each sample was digested and triturated to obtain single-cell or x-nucleus suspensions. Individual cells or nuclei were sorted into 8-well strip PCR tubes by FACS and lysed. SMART-Seq v4 was used to reverse-transcribe and amplify full-length cDNAs. cDNAs were then tagmented by Nextera XT, PCR-amplified, and processed for Illumina sequencing. Initial clustering of sequenced cells was performed independently for each of the species. To further distinguish magno- and parvocellular types the human data was clustered with the macaque data. For cross-species comparison, macaque, human and mouse data were co-clustered using Seurat v3.

### Transcriptomic cell types in macaque dLGN and pulvinar

Single nuclei were isolated from three macaque donors across two species from dLGN and pulvinar. The nuclei were subjected to snRNA-seq, mapped to *Macaca mulatta* genome, clustered with Seurat, and the relationship among clusters was explored in 2D-UMAP projections. This projection revealed heterogeneity within clusters that is driven by donor identity (**Fig. S2A**). Interestingly, differentially expressed genes between donors were related to neuronal signaling and connectivity and not to metabolic or activity-dependent effects (**Table S2**). To explore cell type diversity shared across donors, we used the fastMNN implementation of Mutual Nearest Neighbors (MNN) which allows for more accurate integration of imbalanced datasets than CCA (Haghverdi et al., 2018) (**Fig. S2B**, Methods). By removing donor-specific signatures, we defined 9 shared neuronal types in macaque dLGN (**Fig. 2A,B, Fig. S2C**). We assigned cell type identities based on known marker genes and dissection location (**Fig. 2C**). The neuronal taxonomy has two major branches, GABAergic and glutamatergic (**Fig. 2A**), that further branch into 4 and 5 types, respectively. We identified two distinct koniocellular types (K1 and K2, **Fig. 2A**) that express konio-specific markers *CAMK2A* and *PRKCG* (Murray et al., 2008). The Pulv cluster corresponds to pulvinar relay neurons and expresses *GRIK3* and *LHX2* (**Fig. 2A**) (Jones and Rubenstein, 2004). Finally, magnocellular projection neurons (M) expressed previously reported markers *ABHD17A (FAM108A), BRD4, CRYAB, EEF1A2, IL15RA, KCNA1, NEFM, PPP2R2C,* and *SFRP2* (Murray et al., 2008), and parvocellular projection neurons (P) expressed *FOXP2* (Iwai et al., 2013) and *TCF7L2* (Murray et al., 2008). We also identified many novel M and P markers that are shared across donors (**Fig. 2D**).

**Figure 2.**
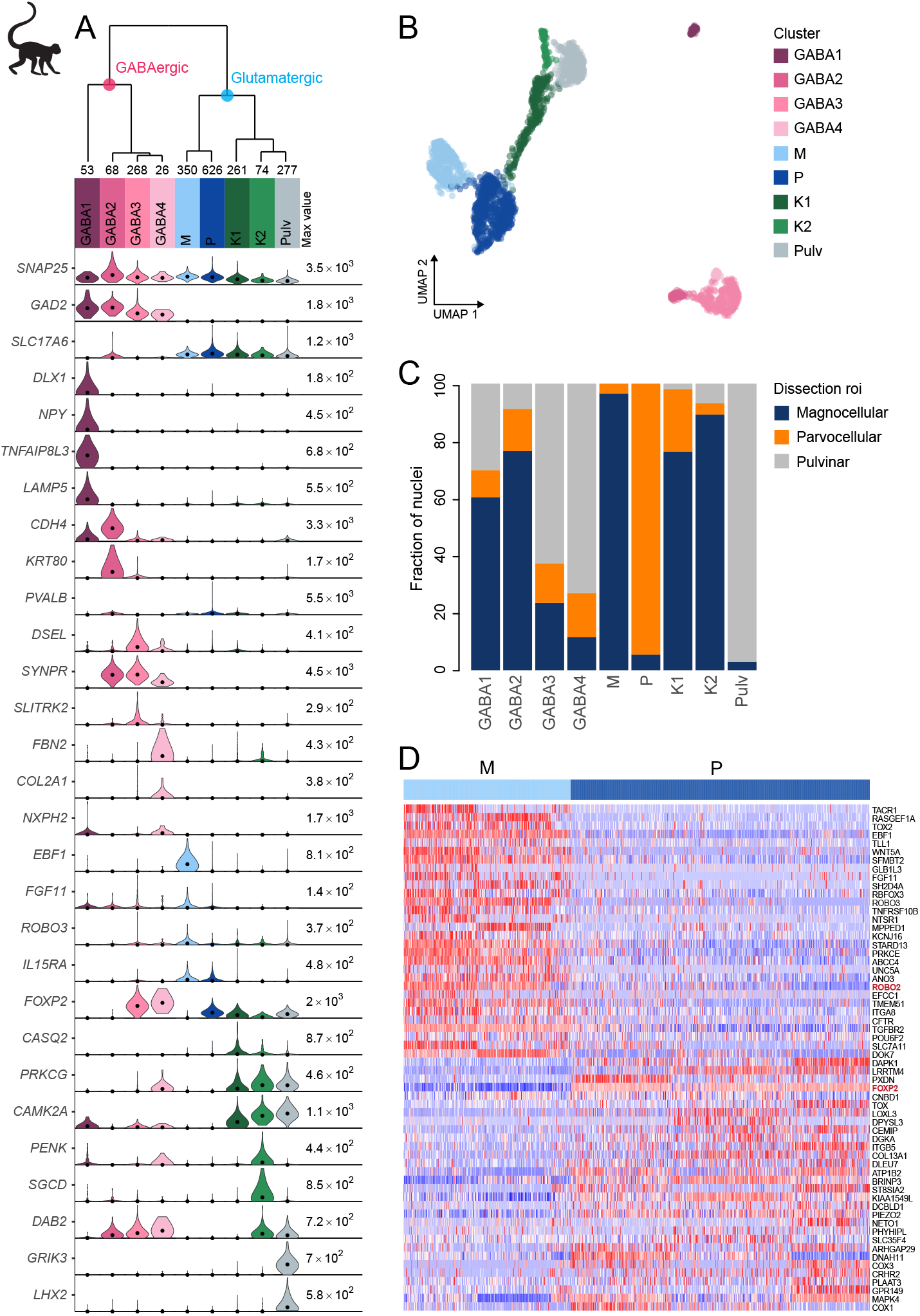
Taxonomy of macaque dLGN and pulvinar neurons by snRNA-seq. **(A)** Hierarchical taxonomy based on median cluster gene expression of 2000 differentially expressed genes in 2003 nuclei from 3 donors across 2 species of macaque. Known marker genes and dissection location were used to assign molecular cluster identity. Numbers of nuclei in each cluster are indicated at the bottom of the dendrogram. **(B)** UMAP representation of macaque dLGN neurons colored by cluster. **(C)** Bar plot representing the fraction of cells derived from a dissection ROI per cluster. **(D)** Heatmap showing the top 60 differentially expressed genes between the M and P clusters.

### Transcriptomic cell types in human dLGN

We define 6 neuronal and 4 non-neuronal types in human dLGN (**Fig. 3A,B; Fig S3A-C**) by transcriptomically profiling individual nuclei isolated from three post-mortem donors as described for macaque above. The neuronal taxonomy has two major branches: GABAergic and glutamatergic, which further branch into 3 types each (**Fig. 3A**). Based on the expression of glutamatergic markers and konio-specific markers (*CALB1*, *CAMK2A* and *PRKCG*, **Fig. 3A**), two koniocellular types could be identified (Hendry and Reid, 2000; Murray et al., 2008). The remaining glutamatergic nuclei belong to a single cluster, to which we assign the parvocellular/magnocellular projection neuron (MP) identity based on the following observations: 1) it is the most numerous glutamatergic type that expresses the known parvocellular/magnocellular marker gene *PVALB* (**Fig. 3A**)(Yan et al., 1996), 2) it does not express koniocellular markers *CALB1*, *CAMK2A* and *PRKCG,* and 3) it contains cells derived from both parvocellular and magnocellular layer dissections (**Fig. 3C**).

**Figure 3.**
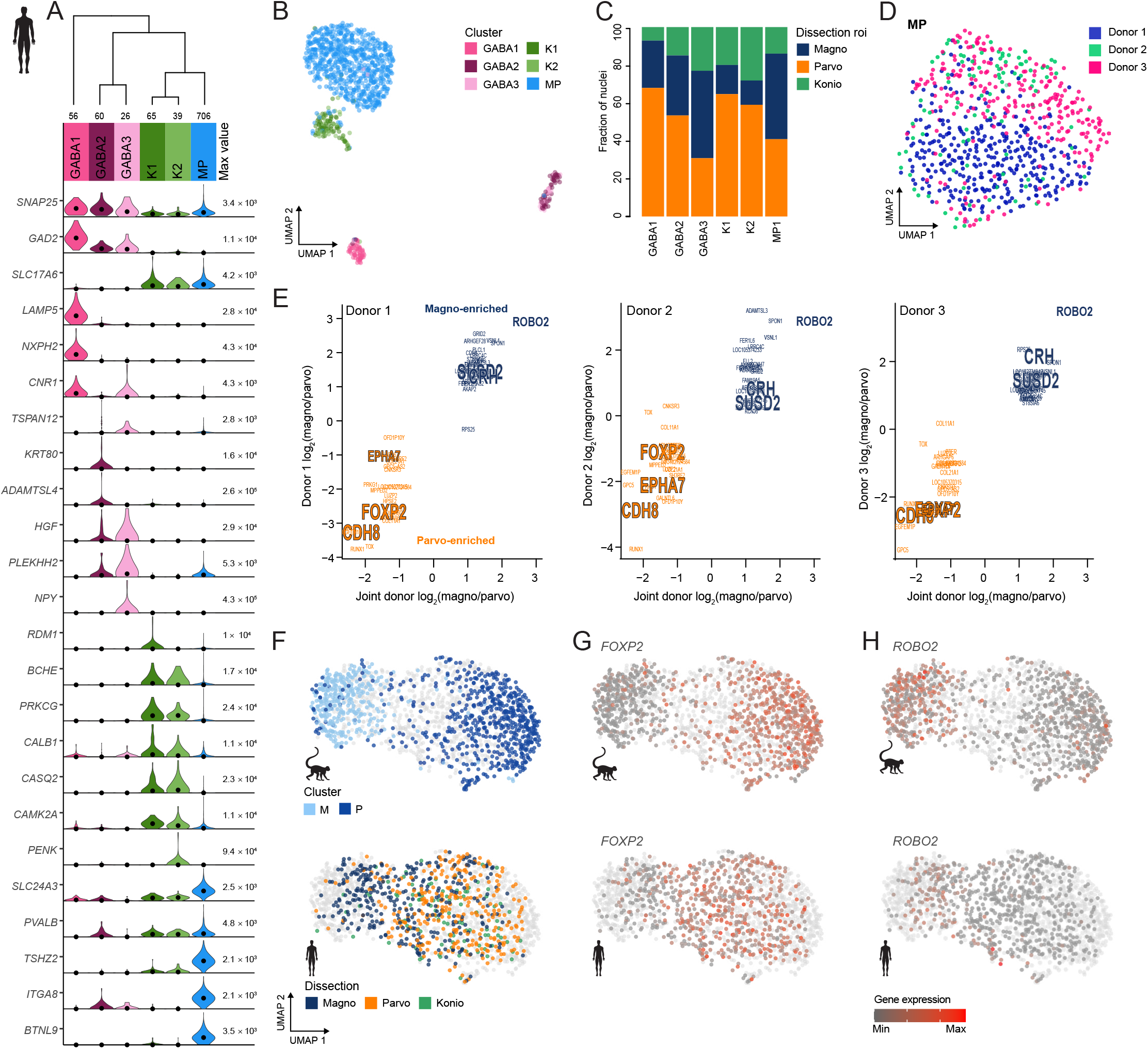
Taxonomy of human dLGN neurons by snRNA-seq. **(A)** Hierarchical taxonomy based on median cluster gene expression of 2000 differentially expressed genes and 946 nuclei from three donors. Known marker genes were used for molecular cluster identity assignment. Numbers of nuclei in each cluster are indicated at the bottom of the dendrogram. **(B)** UMAP representation of human dLGN neurons colored by cluster. **(C)** Bar plot representing fraction of cells derived from a dissection ROI per cluster. **(D)** UMAP representation of neurons within the MP cluster colored by donor. **(E)** 32 Single cell marker genes show consistent enrichments between magno and parvo dissections in three donors. Select marker genes are highlighted. **(F)** UMAP representation of joint analysis of human MP and macaque M and P nuclei using CCA. The macaque nuclei are labeled based on cluster identity defined by single species analysis and the human nuclei within the MP cluster are labeled by dissection ROI. **(G-H)** UMAP representation, as in panel **G**, showing expression of the parvocellular marker *FOXP2* **(G)** or the magnocellular marker *ROBO2* in the macaque and human MP clusters **(H)**.

We detect donor-related heterogeneity in the human MP cluster (**Fig. 3D**). This heterogeneity is unlikely due to sampling different subregions of dLGN across donors because we did not find any significantly differentially expressed genes between anterior and posterior dLGN dissected from individual donors (**Fig. S3D**).

Next, we looked for signatures of M and P types within individual donors. Among the MP nuclei from one donor, we observe a gene expression gradient that corresponds to magno and parvo dissections (**Fig. S3E**). To examine this gene expression gradient in more detail, we analyzed co-clustering diagrams, which define how often nuclei are placed in the same cluster after 100 iterations of clustering. In all three donors, these co-clustering diagrams show substructure within the MP type that corresponds to magno and parvo dissections (**Fig. S3F**). Examination of MP cells within each donor and comparison to joint donor signatures shows consistent gene expression differences between magno- and parvo-enriched genes (**Fig. 3E**). We also confirmed expression of select marker genes by RNA *in situ* hybridization (ISH) (**Fig. S3G,H**), including newly discovered markers for magno (*CRH* and *SUSD2*), parvo (*EPHA7*), or magno and parvo (*BTLN9*) cells.

The gene expression differences between glutamatergic neurons from magnocellular and parvocellular dissections in some of the human donors were subtle, but they were clearly observable in the macaque dataset (**Fig. 2D**, **Fig. S3I**). We therefore wondered if the MP gene expression gradient in human would align with the M and P cluster division in macaque. To investigate this potentially shared gene coexpression pattern between species, we examined human nuclei and batch-corrected macaque nuclei together by employing CCA in Seurat v3. After the cross-species integration, the macaque M and P clusters remain well segregated, whereas the nuclei from the human MP cluster form a continuum. Encouragingly, the human MP continuum aligns with the macaque M and P clusters, that is, the human M-pole nuclei align with the macaque M cluster, and likewise, the human P-pole nuclei align with the macaque P cluster (**Fig. 3F**). This indicates that there is a shared signature between species in the magno/parvo cell populations. *FOXP2* is a robust marker of parvocellular neurons (Iwai et al., 2013) and is enriched in the macaque P cluster and overlapping nuclei in the human MP cluster (**Fig. 3G**) compared to other nuclei. Likewise, *ROBO2* is enriched in the macaque M cluster and overlapping human nuclei (**Fig. 3H**). Despite the differences among donors, species, and nuclear quality, integration of transcriptomic data enabled us to identify a common axis of gene expression variation that is conserved among macaque and human and aligns with the previously defined magno/parvo anatomical axis.

### Transcriptomic cell types in mouse dLGN, LP and LGv

To profile cells from mouse dLGN while examining the reported diversity in relay neurons (Krahe et al., 2011; Cruz-Martin et al., 2014), we performed dissections that enriched for core and shell regions of dLGN. We identified 12 neuronal and 3 non-neuronal types from dLGN dissections in mouse (**Fig. 4A,B; Fig S4A-C**). The 12 neuronal types could be further divided into 9 GABAergic and 3 glutamatergic types. However, due to the small size of dLGN in mice, we suspected that some of these types likely originated from neighboring areas that could not be clearly separated by microdissections. Therefore, as controls, we also profiled single cells from nearby thalamic nuclei, LP and LGv. To assign anatomical location to clusters, we identified differentially expressed genes selective to each cluster and then examined expression patterns of these marker genes in the Allen Brain Atlas RNA in situ hybridization (ISH) data (Lein et al., 2007). Based on these marker genes, we assigned three GABAergic types to dLGN, whereas the remaining GABAergic types were likely from adjacent thalamic nuclei, including LP, LGv, and the reticular thalamic nucleus (RTN) (**Fig. 4C**). Based on the dissection area as well as the differentially expressed genes, all glutamatergic cells from dLGN belong to a single cluster “dLGN” (**Fig. 4A-D**). Cells within this cluster are not homogeneous, with the major axis of gene expression variation corresponding to the core vs. shell anatomical axis (**Fig. 4E**). To confirm this anatomical and gene expression heterogeneity *in situ,* we identified differentially expressed genes between cells isolated from shell and core of dLGN (**Fig. 4F**) and validated expression of a subset of genes by multiplexed RNA ISH (**Fig. 4G**, **Fig. S4D,E**). We confirm our findings that neurons in the shell express higher levels of *Necab1* and *Calb1,* and neurons in the core more highly express *Pvalb* and *Scnn1a.* A small subset of neurons coexpress *Pvalb* and *Necab1,* suggesting that there are ‘intermediate’ cells in dLGN that share shell- and core-like properties. In agreement with this finding, Calb1 protein has been previously reported to be more highly expressed in the dLGN shell compared to the core as measured by immunohistochemical labelling (Grubb and Thompson, 2004).

**Figure 4.**
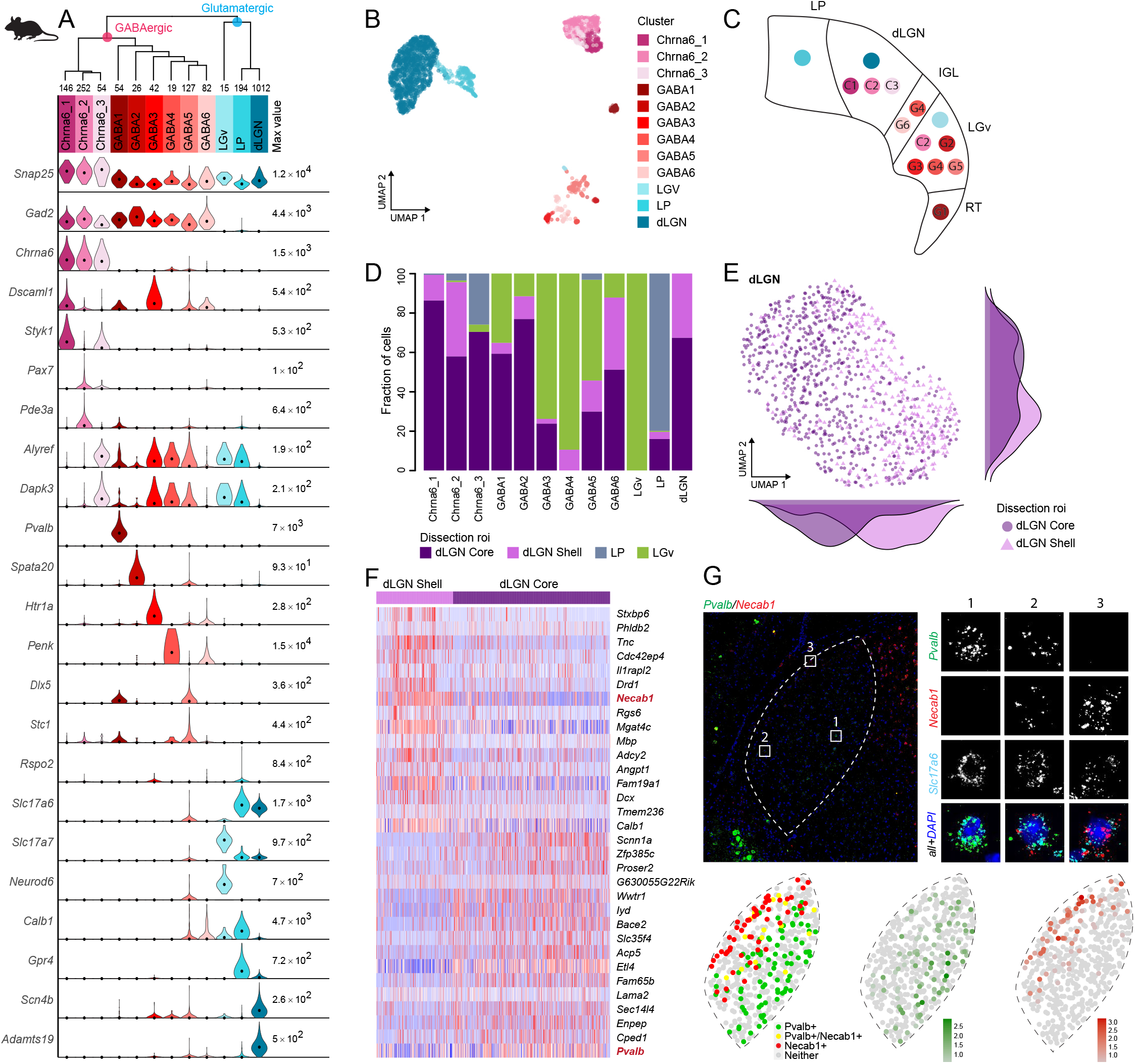
Taxonomy of mouse dLGN and nearby neurons by scRNA-seq. **(A)** Hierarchical taxonomy based on median cluster gene expression of >2000 differentially expressed genes and 2020 cells. Known and newly discovered marker genes were used to assign molecular cluster identity. **(B)** UMAP representation of mouse dLGN neurons colored by cluster. **(C)** Schematic representation of the relevant thalamic nuclei in the mouse brain with colored dots representing cell types identified in this study. Based on cell type specific marker expression and using the Allen Brain Atlas ISH data the anatomical location of cell types could be determined. **(D)** Bar plot representing fraction of cells derived from a dissection ROI per cluster. **(E)** UMAP representation of neurons from the dLGN cluster colored by dissection ROI. The density plot in the margin shows the distribution of cells dissected from mouse dLGN-core (dark purple) and mouse dLGN-shell (light purple) along the x- and y-axis. **(F)** Heatmap showing top 30 differentially expressed genes between cells obtained from dLGN shell and core dissections belonging to dLGN cluster. **(G)** Confirmation of differential expression of *PVALB* and *NECAB1* between shell and core of mouse dLGN by single molecule fluorescence *in situ* hybridization by RNA-scope.

### Cross-species analysis of neuronal cell types in dLGN

To examine cross-species correspondence of the dLGN transcriptomic cell types, we integrated the macaque and human snRNA-seq datasets with the mouse scRNA-seq dataset using CCA in Seurat v3 (**Fig. 5A-C**) for a dataset of n=4,979 neurons. 2D-UMAP projections show extensive intermingling of GABAergic types across the three species but more apparent separation between glutamatergic types (**Fig. 5B,C**). To assess the correspondence, we generated an integrated taxonomy of the cell types (n=17) identified by analysis of each species independently and compared this to the integrated clustering result (n=10 cell types) (**Fig. 5D**). The integrated taxonomy has two major branches: GABAergic and glutamatergic. The integrated GABAergic class contains 16 species-specific types that map to 4 integrated types (**Fig. 5D, Fig. S5A**). Two integrated types are species-specific and likely represent neurons that were sampled from outside of dLGN, including the mouse GABA1 type (likely representing RTN neurons) and the macaque GABA2 type. The mouse GABAergic types can be roughly divided into two groups based on the expression of key transcriptional regulators (*Sox14, Lef1, Otx2, Nkx2-2,* and *Dlx5*) that reflect their developmental origin in the midbrain (*Sox14+*) or forebrain (*Sox14-*) (Scholpp and Lumsden, 2010; Jager et al., 2020). All mouse GABAergic types in dLGN and some types in the ventrolateral geniculate nucleus (LGv) and the intergeniculate leaflet (IGL) express *Sox14,* which is consistent with previous reports (Sellers et al., 2014; Jager et al., 2016). Similarly, two macaque and two human GABAergic types express *SOX14* and are homologous to the *Chrna6/Sox14-expressing* GABAergic dLGN types in mouse. In addition, one macaque and one human cell type are homologous to the mouse GABA2-6 types which are defined by expression of *Npy, Nkx2-2,* and *Dlx1-6* (**Fig. S5B-D**). Interestingly, the mouse GABA2-6 types are found in nuclei adjacent to dLGN and integrate with the macaque GABA1 type and human GABA1 type. The human GABA1 type was validated to be localized to dLGN based on RNA ISH (**Fig. S3H**) and represents ~40% of GABAergic neurons. In summary, we identify two major groups of GABAergic neurons that are conserved across species and correspond to different embryonic origins.

**Figure 5.**
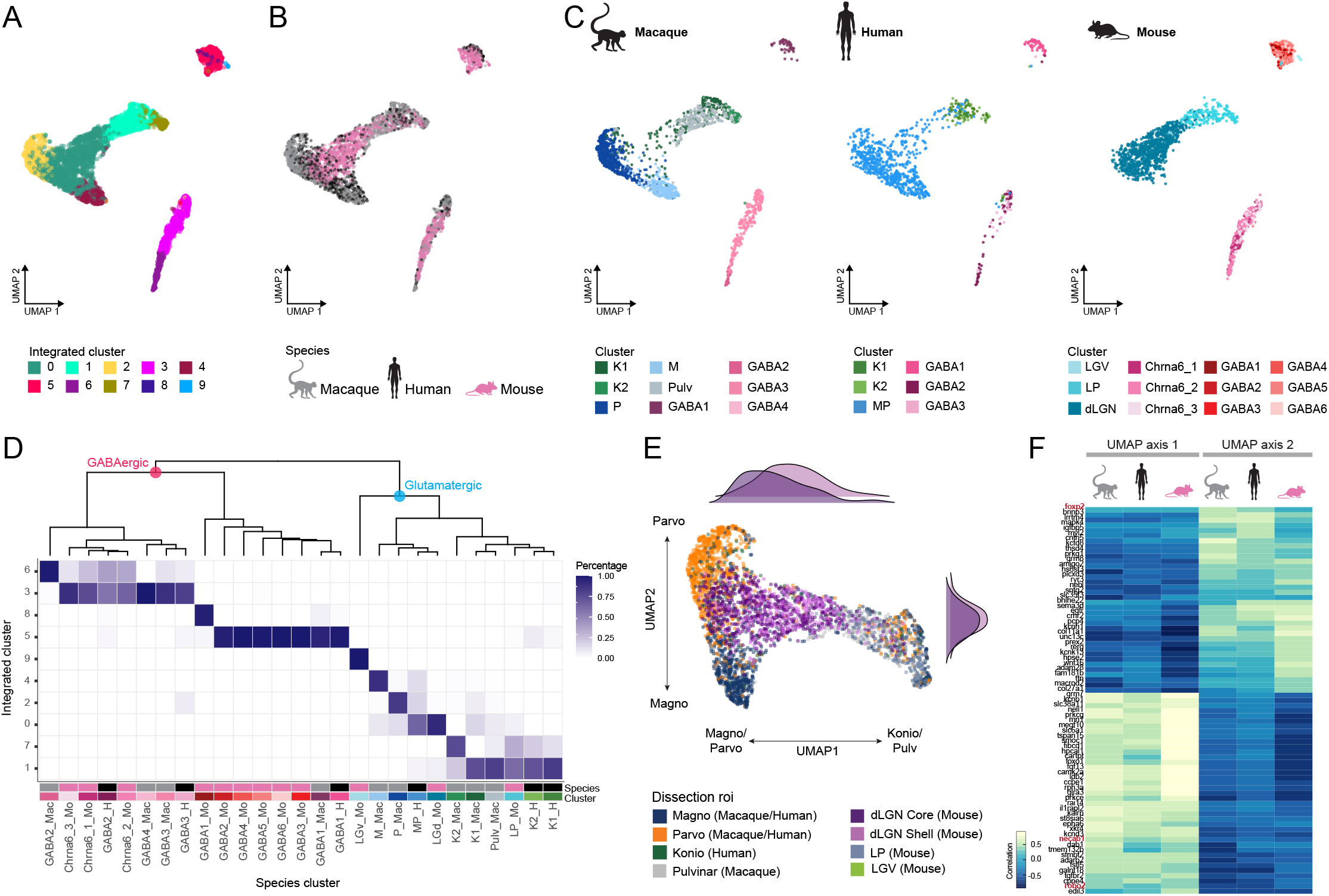
Cross-species integrative analysis. **(A-C)** All neurons from macaque, human, and mouse were integrated using Seurat v3. UMAP representation of the integrative analysis colored by integrated cluster call **(A)**, colored by species **(B)**, or colored by species independent cluster call **(C)**. **(D)** Correspondence of species-specific clustering and integrative clustering. The heatmap illustrates the proportion of the species-specific cell types contributing to the integrated cluster. **(E)** Representation of glutamatergic neurons selected from UMAP as represented in panels A and B colored by dissection ROI. The density plot in the margin shows the distribution of cells dissected from mouse dLGN-core (dark purple) and mouse dLGN-shell (light purple) along the x- and y-axes. **(F)** Heatmap showing the correlation of the gene expression along the UMAP axes represented in panel C per species. For each axis the top 30 genes are shown.

Glutamatergic types did not align as clearly across species as GABAergic types, and mouse types were particularly distinct (**Fig. 5C,D**). Homology between macaque and human types can be resolved better when mouse cells are excluded (**Fig. S5E**). As expected, glutamatergic mouse LGv neurons clustered separately because this region was not sampled in primates. Transcriptomic signatures of mouse LP neurons resembled macaque inferior pulvinar neurons, the homologous structure in primates, and closely related koniocellular neurons. Like the integration of human and macaque data shown in **Fig. 3F**, the human MP cells form a gradient along one axis with the macaque M and P types populating the distinct ends of that same gradient (**Fig. 5C-E**). However, the mouse dLGN cluster variation not only aligns with this gradient of macaque and human magno/parvo types, but rather spans both the continuum between the magno and parvo types along UMAP axis 2 and the continuum between magno/parvo and konio/pulvinar/LP types on UMAP axis 1 (**Fig. 5E**). Intriguingly, more shell-than core-dissected neurons from mouse dLGN resembled konio/pulvinar neurons, consistent with reported similarities in their connectivity (Bickford et al., 2015). For each species, we correlated gene expression with position along these axes and found many genes with graded expression changes along both axes. There is clear conservation in the expression pattern of genes like *ROBO2*, *FOXP2*, and *CAMK2A*, along UMAP axis 1, corresponding to the magno/parvo to konio/pulvinar difference (**Fig. 5F**).

## DISCUSSION

We used unsupervised clustering to define transcriptomic cell types in a well-studied part of the mammalian thalamus, dLGN, and examined the correspondence of these types to previously described morphological, connectional and physiological differences in excitatory relay neurons (Hendry and Reid, 2000; Krahe et al., 2011; Cruz-Martin et al., 2014).

Strong donor-specific molecular signatures were found in both macaque and human datasets. Donor effects may be driven by the greater genetic diversity of primate donors than inbred mice included in this study. It is also possible that some other effects are at play, such as differences in cell state, although we find this possibility unlikely as differentially expressed genes were not enriched for previously identified activity-dependent genes (**Table S2**). It is unlikely that the differences are due to single-nucleus vs. single-cell profiling, as we have previously shown excellent correspondence between taxonomies obtained from single cells and single nuclei for layer 5 cortical neurons in mice (Bakken et al., 2018). The higher consistency among cells, and the resolution of transcriptomic signatures for the macaque compared to human is likely to be of technical nature as we used fresh macaque tissue as opposed to frozen human tissue with 18.5-25 h post-mortem interval. Interestingly, non-human primate donor effects were more prominent among glutamatergic neurons than among GABAergic neurons in LGN and all neurons in human neocortex (Hodge et al., 2019), suggesting that biological variability differs across cell types or cell types vary in their robustness to technical artifacts.

Our data revealed more GABAergic neuron types identified in mouse than in macaque or human. This could be due to sampling 2-fold more GABAergic neurons from the mouse, including from thalamic nuclei adjacent to dLGN, and increased gene detection from mouse whole cells (~8000 genes) versus primate nuclei (~5000 genes). Nevertheless, two major groups of GABAergic types are conserved across the three species (**Fig. 5D**). Two recent studies showed that at least some of the dLGN interneuron developmental programs are conserved between primates and rodents, leading to homology in mature cell types (Golding et al., 2014; Jager et al., 2020). GABAergic neuron diversity in primate dLGN may be increased by an expanded distribution of forebrain-derived, *SOX14*-negative neurons that are found predominantly in medial thalamic nuclei in mouse. Interestingly, this population of neurons has expanded in larger-brained primates, since *SOX14*-negative neurons represent less than 10% of GABAergic cells in marmoset dLGN (Jager et al., 2020), 15% in macaque, and 40% in human.

Consistent with previous results (reviewed in (Hendry and Reid, 2000)), we find that primate koniocellular relay neurons are clearly distinguishable from their magno- and parvocellular counterparts. Moreover, we identify two subtypes of koniocellular neurons that may correspond to two populations reported in macaque that have distinct laminar distributions in dLGN and projection patterns in V1 (Casagrande et al., 2007). We also find that the koniocellular relay neurons are more transcriptomically similar to inferior pulvinar neurons than to magno- or parvocellular neurons. This is consistent with their shared inputs from retina and superior colliculus and cortical projection targets (Huo et al., 2019) and supports a close functional relationship between dLGN and inferior pulvinar. Magno- and parvocellular neurons in both macaque and human are, by comparison, transcriptomically similar to each other. Likewise, we find that relay neurons in mouse dLGN cannot easily be separated into discrete types based on transcriptomic information, although we can detect a primary axis of heterogeneity between core and shell. Similarly, a recent study analyzing single cell transcriptomes across different thalamic nuclei from several projection systems described across and within-nucleus heterogeneity (Phillips et al., 2019). In agreement with our data, among the excitatory neurons from visual thalamus (LP and dLGN), a transcriptomic gradient was observed with the *Pvalb*-expressing projection neurons on one end and *Calb1-* and *Necab1*-expressing neurons on the other, as well as cells co-expressing the three genes in the middle of the gradient.

A recent study assessed gene expression changes across several time points during postnatal development of dLGN using single-cell RNA-sequencing, showing increased transcriptional heterogeneity between postnatal days (P)10 and P16 when compared to later time points in mouse (Kalish et al., 2018). The period between P10 and P14 is a critical period when eye opening occurs and synaptic remodeling in LGN peaks. The RNA-seq results indicate that transcriptional heterogeneity may peak during synaptogenesis and synaptic partner matching (Hooks and Chen, 2006; Iwai et al., 2013). A similar phenomenon has been described in *Drosophila* where the transcriptomes of closely related types of olfactory projection neurons differ the most during circuit assembly but are highly similar at the adult stage (Li et al., 2017). We speculate that maturing relay neurons in dLGN during mouse development may show more discrete transcriptomic signatures than we observe in the adult.

In adult mouse dLGN, three distinct neuronal types, X-, Y-, and W-like, can be identified based on their morphology but have similar electrophysiological properties (Krahe et al., 2011; Bickford et al., 2015). Based on the unsupervised clustering presented in this manuscript, the distinct morphological cells types cannot be identified based on their transcriptomes alone. Likewise, in macaque and human, magno- and parvocellular neurons have highly distinct morphologies, connections, and locations in the dLGN, yet are transcriptomically similar. Other groups have made similar observations where cell types can be clearly distinguished based on morphology and/or electrophysiological properties and not transcriptomic properties and vice versa (Cadwell et al., 2016; Fuzik et al., 2016; Mayer et al., 2019). The discrepancy between the clear morphological, electrophysiological and positional distinctions on the one hand, and the more nuanced transcriptional differences among cell types on the other hand, reinforces the notion that cell type identification is best addressed by a multimodal approach. Indeed, in a recent study, heterogeneity in mouse cerebellar molecular layer interneurons could only be clarified by joint characterization of gene expression, morphology and physiological properties (Kozareva et al., 2020). They observed a continues variation in gene expression in the unipolar brush cells that corresponds to distinct electrophysiological properties, but not to differences in morphology. This study accentuates the need to capture the full range of cell features by obtaining multimodal data. Using multimodal methods like Patch-seq will enable definition of cell types that vary based on their morphology, connectivity, firing patterns and other attributes relevant to their role in neural circuits (Gouwens et al., 2020; Peng et al., 2020). In the future, a clearer definition of discrete and continuous heterogeneity in transcriptomic landscapes may be enabled by new and improved experimental and analytical methods, such as spatial transcriptomics, that can comprehensively sample RNA transcripts from cells *in situ* (Lein et al., 2017).

## ACKOWLEDGEMENTS

We would like to thank the In vivo Sciences team at the Allen Institute for mouse husbandry, the Tissue Procurement, Tissue Processing and Facilities teams for assistance with the transport and processing of postmortem and neurosurgical brain specimens, the Molecular Biology department for processing samples for single cell RNA-sequencing, and the Washington National Primate Research Center for macaque tissue (P51 OD010425). This work was funded by the Allen Institute for Brain Science. The authors thank the Allen Institute founder, Paul G. Allen, for his vision, encouragement and support.

## AUTHOR CONTRIBUTIONS

B.T., G.M., T.E.B., V.M., and H.Z. conceptualized the study. K.A.S. organized and managed the sc/sn RNA-seq pipeline. D.B. performed sc/sn RNA-seq. G.H. provided macaque tissue. R.D.H., T.N.N., N.D., S.P., T.C. and B.P.L. performed single-cell isolation. T.E.B., V.M., C.vV., Z.Y., A.S., J.G., L.T.G. and B. T. performed transcriptome data processing analysis. R.D.H., T.N.N., and E.G. performed RNA *in situ* hybridization with RNAscope. A.B. and J.P. managed transcriptomics pipeline establishment. S.M.S. provided program management support. H.Z. and E.L. led the Cell Types Program at the Allen Institute. C. K. provided funding, institutional support and feedback on the project and manuscript. T.E.B., V.M., C.vV., Z.Y., L.T.G., T.N.N., R.D.H. and B.T. prepared the figures. B.T., T.E.B., V.M., and C.vV. wrote the manuscript in consultation with all authors.

## METHODS

### Overal procedures and data analysis

Full experimental and data processing procedures are available at the Allen Institute web site within a detailed white paper: http://help.brain-map.org/display/celltypes/Documentation?preview=/8323525/10813526/CellTypes_Transcriptomics_Overview.pdf. Below, we list specific aspects that pertain only to this study.

### Mouse breeding and husbandry

All procedures were carried out in accordance with Institutional Animal Care and Use Committee protocols 1508, 1510, and 1511 at the Allen Institute for Brain Science. Animals were provided food and water *ad libitum* and were maintained on a regular 12-h day/night cycle at no more than five adult animals per cage. Animals were maintained on the C57BL/6J background. Experimental animals were heterozygous for the recombinase transgenes and the reporter transgenes. We utilized four Cre lines crossed to the tdT-expressing Cre reporter *Ai14* (Madisen et al., 2010): one panneuronal (*Snap25-IRES2-Cre*) (Harris et al., 2014), one pan-glutamatergic (*Slc17a6-IRES2-Cre)(Vong* et al., 2011), and two pan-GABAergic lines (*Gad2-IRES-Cre* and *Slc32a1-IRES-Cre)(Tong* et al., 2008; Taniguchi et al., 2011).

### Macaque tissue

The brain tissues of adult *Maccaca nemestrina* (southern pig-tailed macaque) and *Macaca fascicularis* were obtained through the Tissue Distribution Program of the Washington National Primate Research Center. All procedures were approved by the Institutional Animal Care and Use Committee of the University of Washington.

### Human tissue

Postmortem adult human brain tissue was collected after obtaining permission from decedent next-of kin. Postmortem tissue collection was performed in accordance with the provisions of the United States Uniform Anatomical Gift Act of 2006 described in the California Health and Safety Code section 7150 (effective 1/1/2008) and other applicable state and federal laws and regulations. The Western Institutional Review Board reviewed tissue collection processes and determined that they did not constitute human subjects research requiring institutional review board (IRB) review. In general, three to five slices were sufficient to capture the targeted region of interest, allowing for expression analysis along the anterior/posterior axis.

### Single cell/nucleus processing for sc/snRNA-seq

We used previously described procedures to perform single-cell and single-nucleus RNA-seq (Bakken et al., 2018; Tasic et al., 2018). In brief, cells and nuclei were isolated by FACS: macaque and human nuclei were stained with the neuronal marker NeuN and NeuN+ nuclei were sorted, whereas mouse cells were collected from several transgenic Cre-driver lines that preferentially label neuronal cells. We reverse transcribed mRNA and amplified cDNA using Smart-seq V4 (Clontech), prepared sequencing libraries using Nextera XT (Illumina), and sequenced the libraries using HiSeq2500 (Illumina). We employed previously described quality control (QC) steps (Bakken et al., 2018; Tasic et al., 2018) to arrive to the final datasets (**Fig. S1**).

### Data processing and analysis

Processing of sequencing data was performed as described before (Bakken et al., 2018; Tasic et al., 2018; Hodge et al., 2019; Bakken et al., 2020). For mouse, raw read (fastq) files were aligned to the mm10 mouse genome sequence (Genome Reference Consortium, 2011) with the RefSeq transcriptome version GRCm38.p3 (current as of 01/15/2016) and updated by removing duplicate Entrez gene entries from the gtf reference file. For human, raw read files were aligned to the GRCh38 human genome sequence (Genome Reference Consortium, 2011) with the RefSeq transcriptome version GRCh38.p2 (current as of 4/13/2015) and likewise updated by removing duplicate Entrez gene entries from the gtf reference file. For analysis of transcriptomes of *Macaca nemestrina* (southern pigtailed macaque), and *Macaca fascicularis* which are the species of macaque used for experiments, we used the genome assembly and annotation of *Macaca mulatta* (rhesus macaque) from the University of Nebraska Non-human Primate Genome center (MacaM_v7.8.2, released 07/17/2016). Alignment to the genome was performed using STAR v2.5.3 (Dobin et al., 2013). Only uniquely aligned reads were used for gene quantification. The observed bimodal pattern in reads per cell for some of the mouse clusters is due to a higher rate of multiplexing, leading to lower sequencing depth, for one batch of cells (**Fig. S1B**). This lower read depth, however, does not result in lower number of genes detected in these mouse cells (**Fig. S1C**). Cells that met any one of the following criteria were removed from the dataset: <100,000 total reads, <1,000 detected genes (counts per million > 0), < 75% of reads aligned to genome, CG dinucleotide odds ratio > 0.5, or doublet score > 0.25. Cells that passed quality control criteria were included in clustering analysis, which was performed using Seurat (Butler et al., 2018; Stuart et al., 2019). The fastMNN implementation of the Mutual Nearest Neighbors method was used to correct for donor effects observed in the macaque dataset. Both CCA and fastMNN attempt PCA subspace alignment. The PCA axes with the highest variance can be lost when using CCA. In cases where cell types in different batches are extremely imbalanced, as is the case for macaque where neurons from pulvinar were collected from one donor, it might lead to incorrect alignment. The MNN procedure uses a different approach. It finds the nearest neighbors across batches. The difference between the paired cells is then used to infer the magnitude and direction of the batch effect across all PCA subspaces to correct the data. Clustering results from individual species were integrated by employing CCA in Seurat v3 (Butler et al., 2018; Stuart et al., 2019).

### RNA *in situ* hybridization

Single molecule RNA *in situ* hybridization by RNAscope (Advanced Cell Diagnostics, Newark, CA) was performed as previously described (Tasic et al., 2018) using fluorescent kits for mouse tissue, and the duplex chromogenic kit for human tissue according to manufacturer’s instructions.

## SUPPLEMENTARY DATA

**Supplementary Table 1. Specimen annotations.** All specimens used in this study are listed with associated clustering, donor, and dissection annotations.

**Supplementary Table 2. Donor specific gene signatures in magno-vs. parvocellular neurons.** Result of differential gene expression analysis of magnocellular vs. parvocellular neurons in macaque and human dLGN per donor.

## SUPPLEMENTARY FIGURE LEGENDS

**Supplementary Figure 1.**
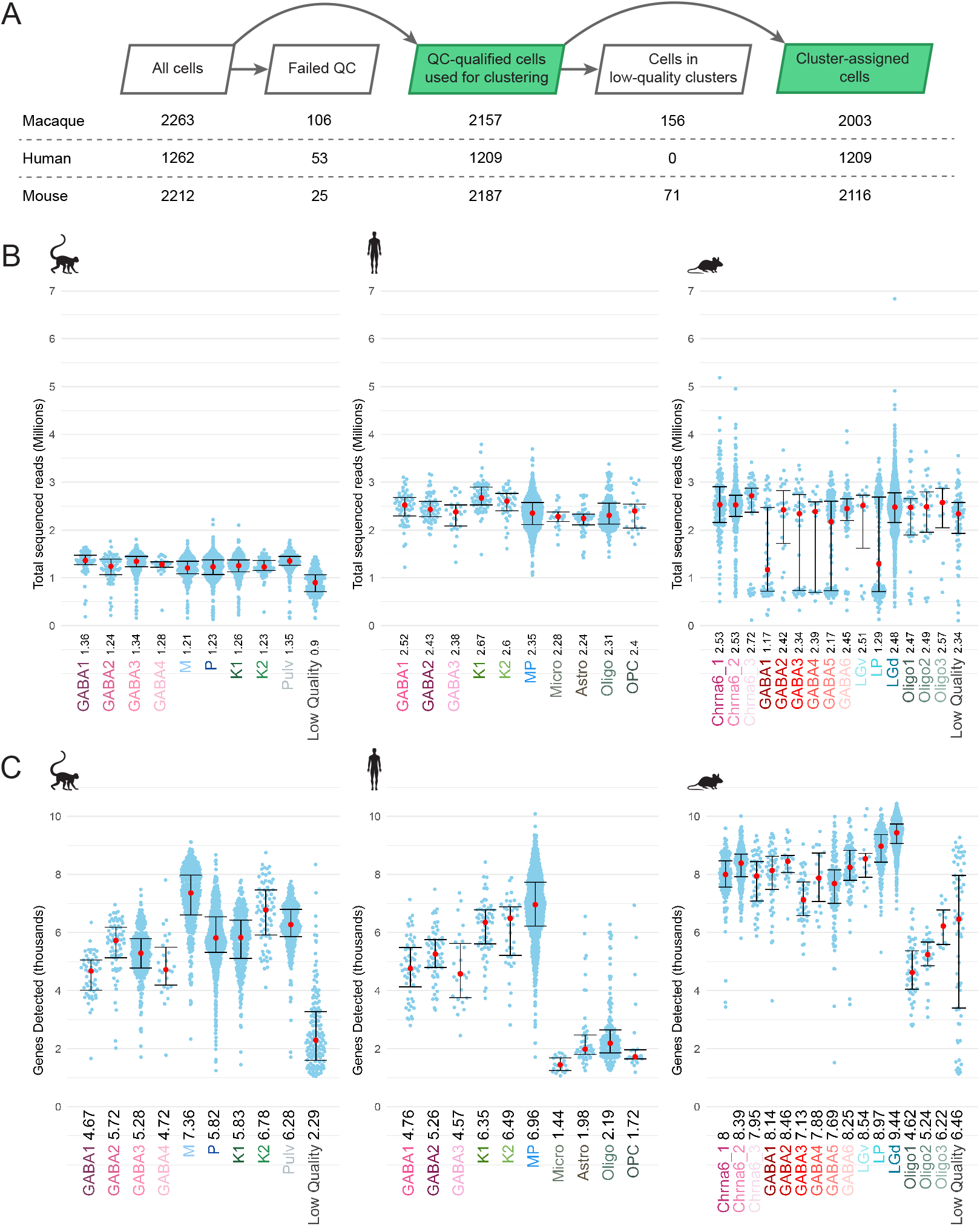
Total reads and gene detection rates in the datasets. Sequencing depth and gene detection for each of the datasets grouped by transcriptomic type. (A) Sequencing depth for all cluster-assigned cells, grouped by transcriptomic cell type per species. Median values in millions of reads are noted adjacent to the cell type label and plotted as red dots; whiskers denote 25^th^ and 75^th^ percentiles. (B) The number of detected genes grouped by transcriptomic type for each species. Median values in thousands of genes are noted adjacent to the cell type label and plotted as red dots; whiskers denote 25^th^ and 75 ^th^ percentiles. (C) Confusion matrix showing cluster membership validation for each species. Validation of cluster membership was performed as described previously (Tasic et al., 2016).

**Supplementary Figure 2.**
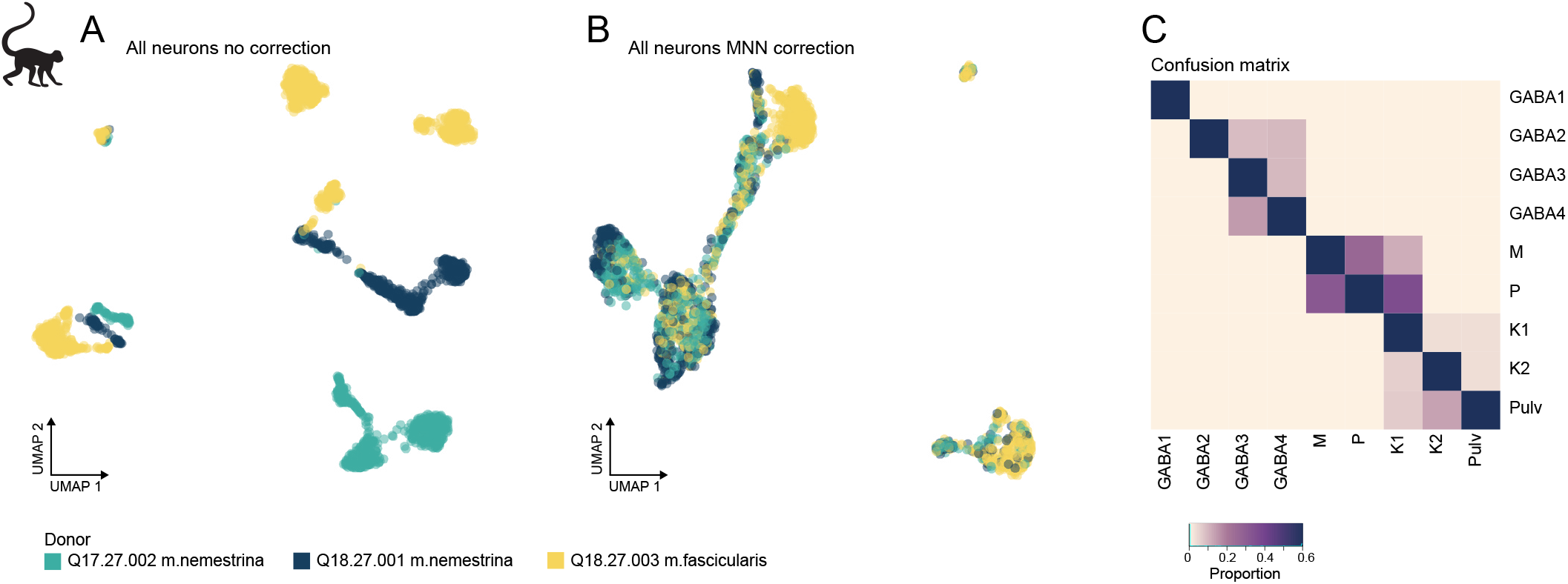
Donor correction in macaque dLGN cells. **(A-B)** UMAP representation of 2003 macaque dLGN neurons colored by donor without correction **(A)** and after fastMNN correction of donor effects **(B)**. **(C)** Confusion matrix showing cluster membership validation for the 9 macaque cell types. Validation of cluster membership was performed as described previously (Tasic et al., 2016)

**Supplementary Figure 3.**
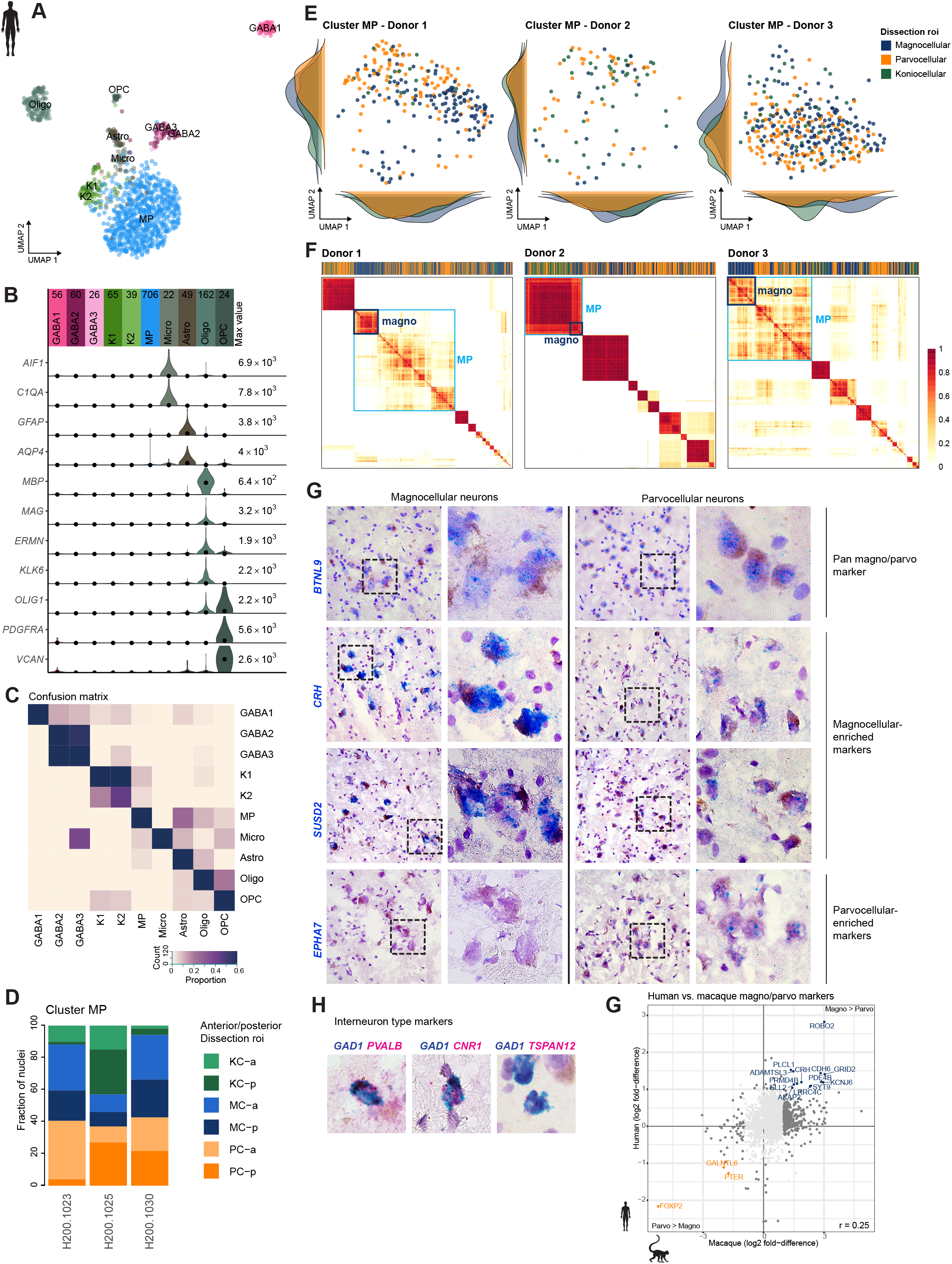
Marker gene expression in magnocellular and parvocellular neurons in Human dLGN. **(A)** UMAP representation of all human dLGN nuclei, neuronal and non-neuronal nuclei, colored by cluster. **(B)** Marker genes expressed in non-neuronal cell types identified in human dLGN. **(C)** Confusion matrix showing cluster membership validation for the 10 human cell types. **(D)** Distribution of dissection ROI per donor within the MP cluster. KC, koniocellular; MC, magnocellular; PC, parvocellular; a, anterior; p, posterior. **(E)** The MP cluster UMAP shown in **Fig 3D** was split by donor and colored by dissection ROI. **(F)** Co-clustering diagrams for three independent donors depicting magnocellular nuclei enrichment within the MP cluster. **(G)** Scatter plot showing consistent single cell marker gene expression between magno- and parvocellular dissections in macaque and human. Select marker genes are highlighted. **(H-I)** Confirmation of select marker gene expression in human dLGN by chromogenic single molecule RNA *in situ* hybridization by RNA-scope. Representative images showing expression of magnocellular and parvocellular marker genes **(G)**. Representative images are shown showing the pan-GABAergic marker GAD1 and specific cluster markers **(H)**.

**Supplementary Figure 4.**
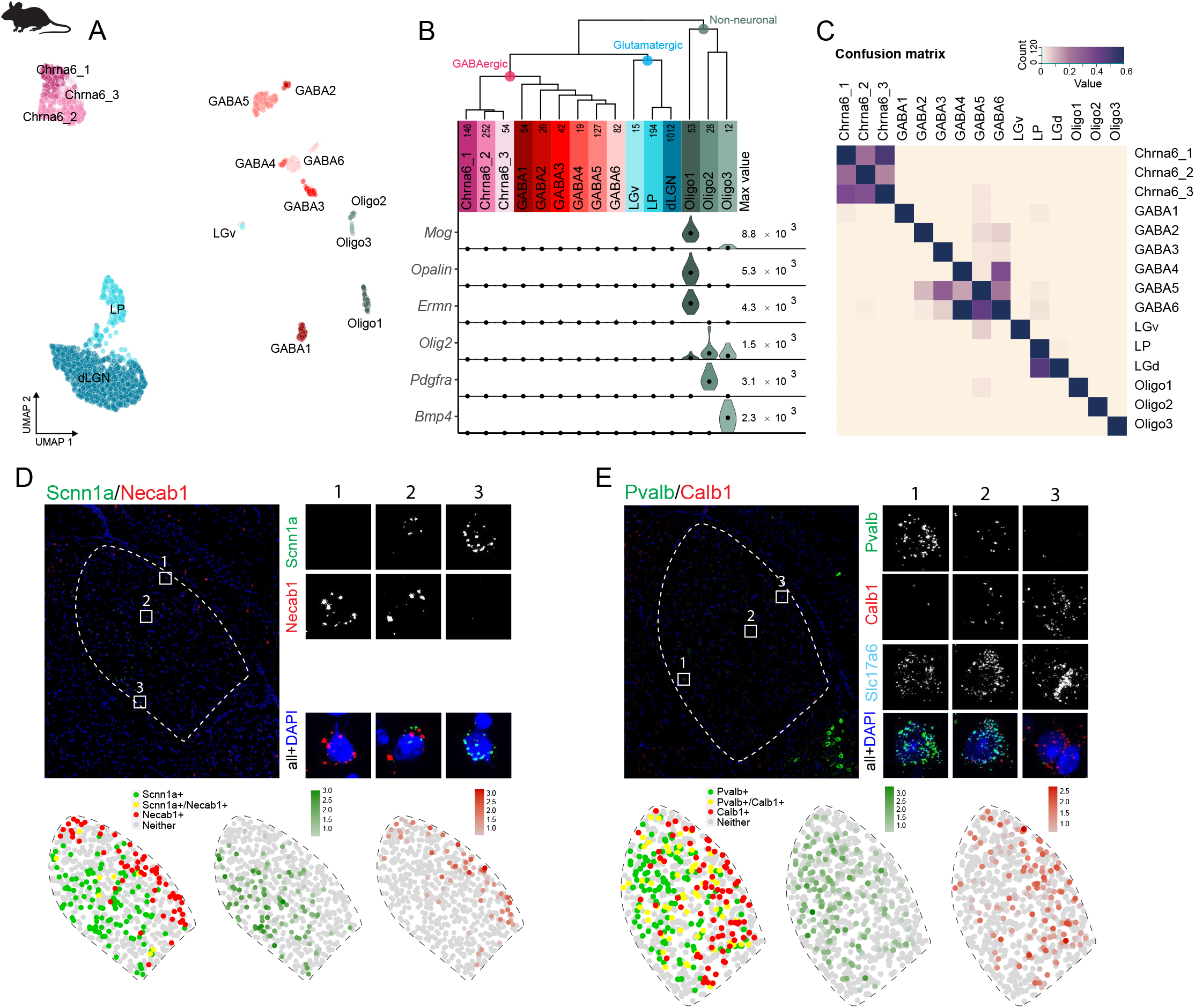
Marker gene expression in mouse dLGN. **(A)** UMAP representation of all mouse dLGN nuclei, neuronal and non-neuronal nuclei, colored by cluster. **(B)** Marker genes expressed in non-neuronal cell types identified in mouse dLGN. **(C)** Confusion matrix showing cluster membership validation for the 15 mouse cell types. **(D-E)** Confirmation of select differential marker gene expression between shell and core of mouse dLGN by single molecule fluorescence *in situ* hybridization by RNA-scope. ISH result showing spatially restricted expression of *SCNNA1* and *NECAB1* **(D)** and expression of *PVALB* and *CALB1* in mouse dLGN **(E)**.

**Supplementary Figure 5.**
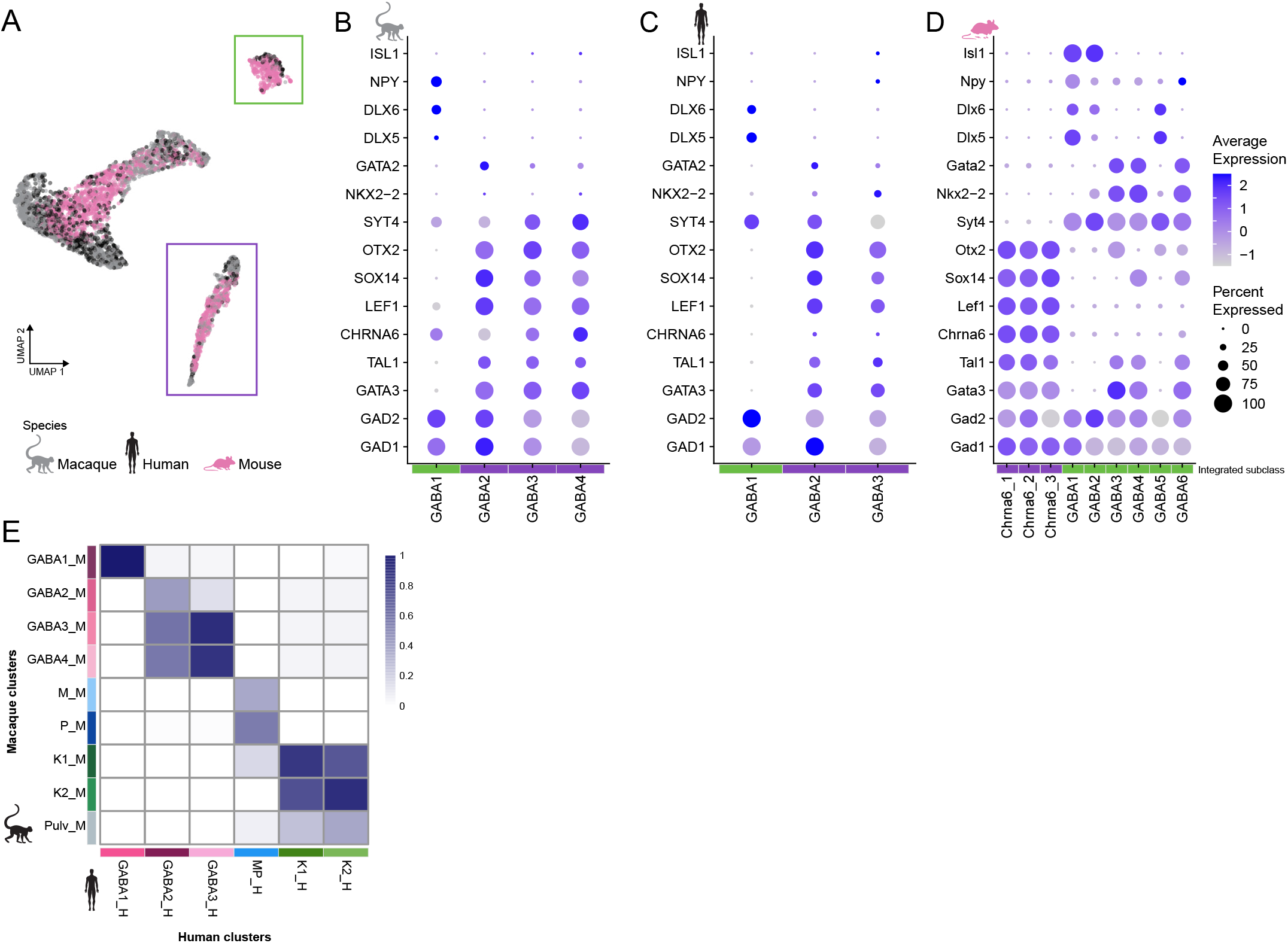
Cross-species comparison of GABAergic types. **(A)** UMAP representation of the integrative analysis. The GABAergic types are colored by species and the glutamatergic types are visible in gray. The GABAergic types are divided in 2 subclasses (denoted by green and purple box) containing cell types from all species. **(B-D)** Dotplot showing expression of key transcriptional regulators of the GABAergic lineage in Macaque **(B)**, human **(C)**, and mouse **(D)** GABAergic types. The color intensity of the dots represents the average expression level, whereas the size of the dot represents the proportion of cells expressing the gene. **(E)** Integration of macaque and human neuronal clusters. The heatmap shows the proportion of cells within-species clusters that align with the integrated cross-species cluster.

## Notes

### Competing Interest Statement

The authors have declared no competing interest.

### Summary of Updates

Proofing update

